# DDTRN: Predicting Bacterial Transcriptional Regulatory Networks Based on Gene Sequences using Dual Descriptor

**DOI:** 10.64898/2026.06.30.735580

**Authors:** Pu Nie, Bin-Guang Ma

## Abstract

Accurate computational reconstruction of bacterial transcriptional regulatory network (TRN) from sequence information alone remains a fundamental challenge in systems biology, particularly for non-model organisms lacking extensive transcriptomic data. We present DDTRN, a sequence-driven framework that formulates TRN inference as a binary classification task over concatenated regulator-target gene sequence pairs and employs a Dual Descriptor (DD) model to predict regulatory interactions. The DD architecture represents a sequence into two learnable components: Composition Weight Map (CWM) and Position Weight Function (PWF). We comprehensively evaluate DDTRN against six conventional machine learning baselines across eight benchmark bacterial datasets, including *E. coli* (DREAM5, RegulonDB), *B. subtilis, S. enterica, C. glutamicum, M. tuberculosis, P. aeruginosa*, and *S. coelicolor*. DDTRN achieves superior overall performance, attaining average AUROC and AUPR scores of 0.869 and 0.868, respectively, with particularly pronounced advantages at lower descriptor ranks where positional weighting compensates for limited sequence context. Systematic sensitivity analyses of rank, embedding dimension, and basis function count reveal stable optimal operating regimes, while subsampling experiments demonstrate strong robustness even with limited training data. Interpretability analyses show that PWF learns distinct periodic contributions across different rank granularities and that CWM preferentially weights meaningful *k*-mers. A case study on *E. coli* dataset further illustrates that DDTRN identifies method-specific candidate targets complementary to those proposed by conventional approaches. By operating solely on genomic sequence, DDTRN provides a scalable, interpretable, and data-efficient framework for bacterial TRN inference in species where expression data are scarce, and it establishes a foundation for future multimodal integration with condition-specific regulatory information.

## 1. Introduction

Transcriptional regulation is a fundamental process in cellular life, crucial for maintaining normal cell growth and function [1]. This process modulates transcriptional efficiency by regulating the activity of RNA polymerase or its associated transcription factor (TF) proteins, thereby influencing gene expression levels and involving virtually every basic activity of life [2]. The transcriptional regulatory network (TRN), which delineates the regulatory relationships between transcription factors (TFs) and target genes (TGs), is often regarded as a functionally upstream regulatory core because it directly controls the initiation and intensity of gene expression [3-5]—a complex, multi-step process that converts specific DNA sequences into functional proteins through precisely regulated stages including transcription, RNA processing, and translation [6]. In the early stages of genomics, reconstructing genome-scale TRN was a highly challenging systems engineering task, with the core difficulty lying in the heavy reliance on massive experimental data to identify transcription factor binding sites (TFBS) and characterize their regulatory activity [7]. However, with the advent and rapid development of high-throughput sequencing technologies in the post-genomic era [8], it has become feasible to readily acquire genomic sequence data and gene expression profiles. Concurrently, continuous advancements in deep learning have opened new technical pathways for constructing high-precision TRNs [9].

Early TRN construction primarily relied on biological experiments such as ChIP-seq to validate regulatory relationships between TFs and TGs on a case-by-case basis. These techniques enabled genome-scale mapping of TFBS, laying a crucial foundation for building large-scale regulatory datasets and subsequent computational modeling [10]. Furthermore, *in vitro* high-throughput assays like DAP-seq, owing to their speed, low cost, and scalability, have provided important means for large-scale TFBS acquisition [11]. On the other hand, when large-scale gene expression data are available, predicting TRNs using RNA-seq data as the core input has become a mainstream approach, leading to the development of a series of computational methods. For instance, GRADIS employs Support Vector Machine (SVM) to reconstruct TRNs based on graph representations of transcriptomic data [12]; GRGNN formulates TRN prediction as a graph classification problem, utilizing Graph Neural Networks for end-to-end learning [13]; the recently proposed PGBTR is a general deep learning framework for bacterial TRN prediction that integrates expression data and genomic information [14]. However, these methods typically depend heavily on large-scale transcriptomic data, which are often plagued by noise, batch effects, and heterogeneity. More critically, for many non-model organisms, it is usually difficult to obtain sufficient quantity or condition-specific RNA-seq data, rendering TRN inference methods that rely solely on expression data less effective in data-scarce scenarios [12-14].

Another core aspect of regulatory prediction is the modeling and characterization of TFBS and their sequence motifs. Traditional methods often use position weight matrices to represent sequence specificity, but their expressive power is limited. Deep learning enables end-to-end learning from raw DNA sequences, significantly enhancing predictive capabilities: DeepBind uses Convolutional Neural Networks (CNNs) to learn the sequence-binding specificity of proteins [15]; DeepSEA takes sequences as input to simultaneously predict multiple regulatory phenotypes, demonstrating the feasibility of sequence-based prediction [16]; DanQ combines CNNs and Bidirectional Long Short-Term Memory networks to model motif features and long-range dependencies [17]; Enformer further integrates long-range genomic interaction information to improve prediction performance [18]. These advances indicate that sequences contain rich regulatory information, making it possible to predict regulatory relationships using sequence as the primary input even in the absence of expression data [15-18]. Therefore, how to construct a model capable of stably predicting genuine regulatory relationships relying solely on widely available genomic sequence information remains a worthwhile research challenge.

Building upon these challenges and opportunities, this work proposes a novel TRN prediction method based on the Dual Descriptor (DD) model [19], termed DDTRN. This method can predict regulatory relationships between TFs and TGs using only gene sequence information. We evaluated DDTRN against other machine learning methods across eight benchmark datasets. Overall, DDTRN achieved optimal performance on most datasets. Moreover, compared to traditional machine learning models, DDTRN offers good interpretability: the model decomposes sequence information into compositional and arrangement information, characterized by Composition Weight Map (CWM) and Position Weight Function (PWF), respectively, thereby providing deep insight into the mechanism of sequence representation related to transcriptional regulation.

## 2. Materials and methods

### 2.1 Benchmark datasets

DDTRN was evaluated on eight bacterial TRN datasets, including the *E. coli* dataset from the DREAM5 challenge [20], two datasets constructed by Gu (14) (RegulonDB_Ecoli, Subtiwiki_Bsubtilis), one dataset from iModulonDB 2.0 [21] (*Salmonella enterica*), and four datasets from Abasy Atlas [22] (*Corynebacterium glutamicum, Mycobacterium tuberculosis, Pseudomonas aeruginosa, Streptomyces coelicolor*). The gene sequences for these bacterial species corresponding to the eight TRN datasets were extracted from reference genomes obtained from NCBI Datasets [23]. **Table 1** displays detailed information for the datasets used in this study.

**Table 1.** Information on the datasets used in this study.

| Dataset | Gene No. | Regulation No. | TF No. |
| --- | --- | --- | --- |
| Dream5_Ecoli | 4297 | 2066 | 334 |
| RegulonDB_Ecoli | 4270 | 5316 | 220 |
| Subtiwiki_Bsubtilis | 4325 | 3033 | 185 |
| *Salmonella enterica* | 2068 | 1808 | 75 |
| *Corynebacterium glutamicum* | 2228 | 3013 | 71 |
| *Mycobacterium tuberculosis* | 3056 | 9813 | 179 |
| *Pseudomonas aeruginosa* | 2627 | 4390 | 91 |
| *Streptomyces coelicolor* | 4863 | 5024 | 160 |

### 2.2 DDTRN model architecture based on Dual Descriptor (DD)

DDTRN was designed as a sequence-driven framework for predicting bacterial transcriptional regulatory relationships from genomic sequences (**Figure 1**). Based on the benchmark datasets, the DNA sequences of the genes with regulatory relationships in the bacterial TRNs were prepared as the input data. For each candidate TF–TG pair, the DNA sequence of the TF gene and that of the target gene were concatenated into a single composite sequence (**Figure 1A**), which was then represented by the vector DD model (tensor form), termed DualDescriptorTS [24]. The final task was formulated as a binary classification problem, where each composite sequence was assigned to either the regulatory or non-regulatory class.

**Figure 1.**
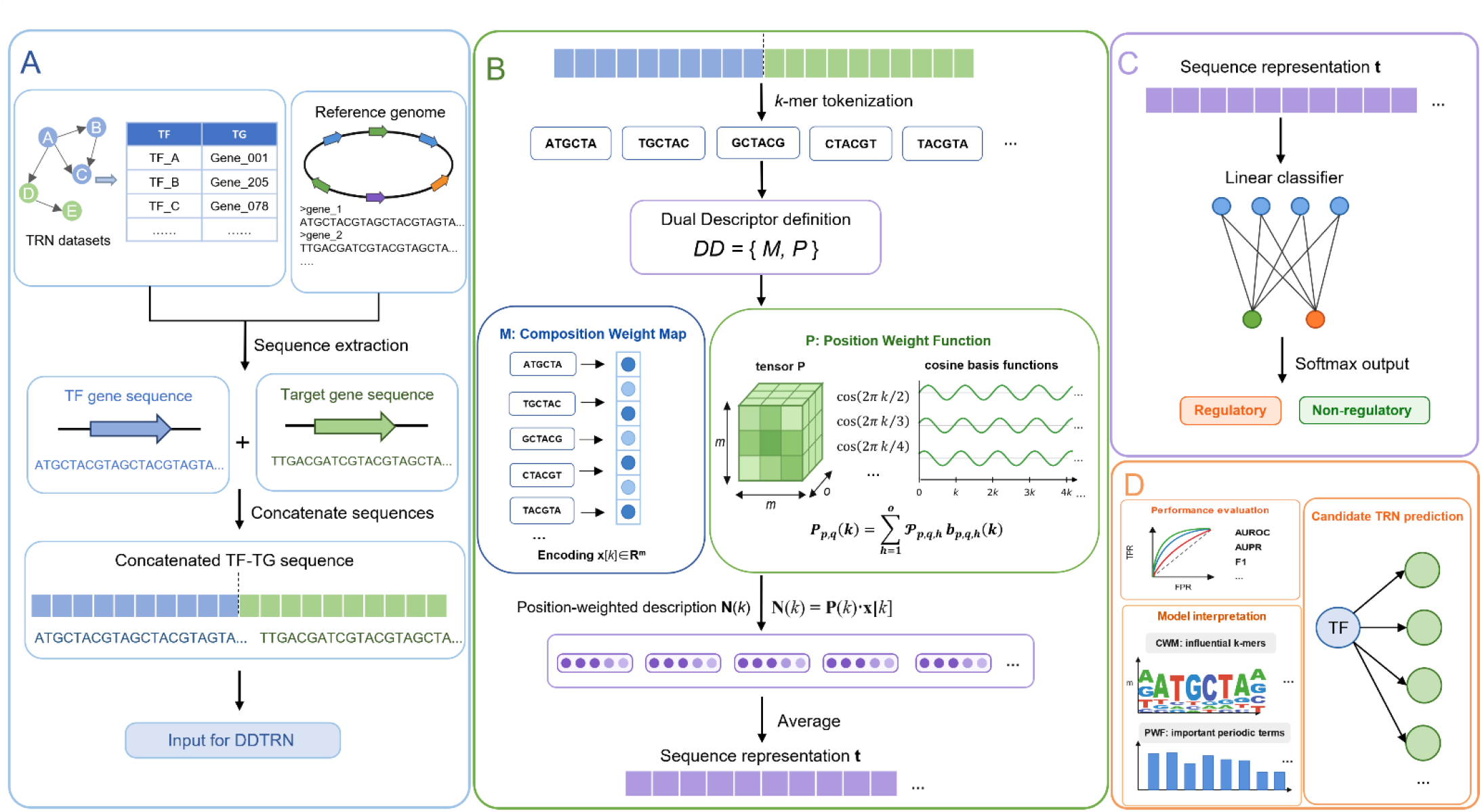
The workflow of DDTRN. (A) Construction of concatenated gene sequences from TF and TG genes. (B) Definition of Dual Descriptor (DD) for sequence representation. (C) Regulatory relationship prediction. (D) Model evaluation and interpretation.

The core of DDTRN is the DD representation: *DD* = {*M, P*}, which decomposes sequence information into two complementary components: the Composition Weight Map (CWM, namely *M*) and the Position Weight Function (PWF, namely *P*) [25]. The CWM describes the compositional contribution of sequence tokens, while the PWF describes their positional or arrangement-dependent contribution (**Figure 1B**). In this way, DDTRN differs from conventional *k*-mer frequency-based representations, which mainly encode how often each *k*-mer occurs but do not explicitly model where and how these sequence fragments are arranged.

Given an input sequence, DD first tokenizes it into rank-length nucleotide fragments. When the descriptor rank is *r*, each token corresponds to an *r*-mer over the alphabet {A, C, G, T}. In the default linear mode, tokens are extracted using a sliding window with stride 1, allowing overlapping *r*-mers to preserve fine-grained local sequence patterns. Each valid *r*-mer token is then mapped by the CWM to a learnable embedding vector **x**[*k*] ∈*R*^*m*^, where *m* is the encoding vector dimension and *k* denotes the token position in the sequence.

To incorporate positional information, DD uses a learnable position-weight tensor **P**∈*R*^*m*×*m*×*o*^, where *o* is the number of basis functions used to expand the PWF. For each token position *k*, a set of cosine basis functions is computed as:

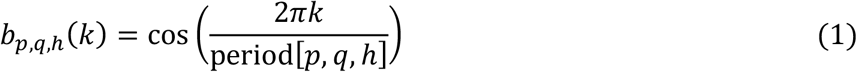

where period[*p, q, h*] is the indexed period, with *p, q, h* indicating the indices in the three dimensions of tensor **P**. In the implementation, the period is defined as:

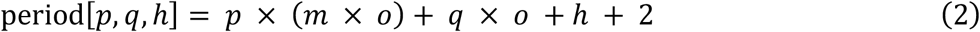

The position-weighted description vector **N**(*k*) for the *k*-th token is then calculated by combining the token embedding, the position weight tensor, and the corresponding cosine basis functions:

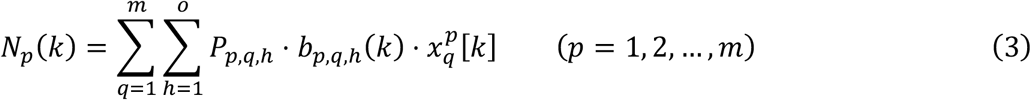

Thus, each **N**(*k*) vector jointly reflects the identity of the local sequence fragment and its position-dependent contribution within the composite TF–TG sequence. The representation of the whole sequence is obtained by averaging all position-level description vectors:

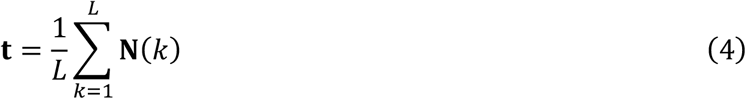

where *L* is the number of extracted tokens. This average description vector **t** serves as the latent representation of the candidate TF–TG pair.

For regulatory relationship prediction, a linear classification head was attached to the DD representation (**Figure 1C**). The classifier maps **t** to two output logits corresponding to the regulatory and non-regulatory classes, and the probability of a regulatory interaction is obtained using the softmax function. During training, the parameters of the CWM, the PWF tensor, and the classification head were optimized jointly by minimizing the cross-entropy loss. Therefore, DDTRN learns sequence representations in an end-to-end manner while retaining the interpretability of the DD framework: the CWM parameters indicate which *r*-mer tokens contribute strongly to the prediction, whereas the PWF tensor reflects which positional or periodic components are important for encoding regulatory sequence patterns.

In this study, the default DD configuration was set to *rank* = 6, *rank_mode* = “drop”, encoding vector dimension *m* = 10, number of basis functions *o* = 10, and description *mode* = linear/nonlinear. These values were selected based on the parameter sensitivity analyses described below. The model was implemented in PyTorch and trained on a GPU when available [24].

### 2.3 Baseline methods

For the baseline methods, six classifiers were employed with their standard implementations: Support Vector Machine (SVM) [26] with radial basis function kernel via SVC(probability=True), Random Forest (RF) [27] with max_features=‘sqrt’, Gradient Boosting Decision Trees (GBDT) [28] with early stopping (n_iter_no_change=10, validation_fraction=0.1), Light Gradient Boosting Machine (LGBM) [29] with learning_rate=0.5, XGBoost [30] with early_stopping_rounds=10, Multilayer Perceptron (MLP) [31] with one hidden layer of 100 units, max_iter=500, and early stopping. The same concatenated TF-TG gene sequences were transformed into fixed-dimensional feature vectors based on *k*-mer frequency counts. Specifically, all possible *k*-mers of length rank were enumerated from the alphabet {A, C, G, T} to form a vocabulary of size 4^rank^. For each sequence, a sliding window of length rank (stride = 1, i.e., linear mode) was used to extract all overlapping *k*-mers; only those fully composed of A, C, G, or T were retained. The occurrence count of each *k*-mer was then normalized by the total number of valid *k*-mers in the sequence, producing a frequency vector of dimension 4^rank^. For SVM and MLP, a standard scaler was applied as a preprocessing step within a pipeline to normalize the *k*-mer frequency features. All models used the same random seed for reproducibility.

### 2.4 Training and testing strategies

A balanced dataset was constructed for each bacterial species by pairing all known positive regulatory interactions (TF–TG) with an equal number of negative samples. Negative samples were generated using a TF-based strategy: for each TF, a target gene was randomly selected from the set of all genes that are neither the TF itself nor a known target of that TF, ensuring that the negative pair did not appear in the positive set. The entire set of labeled pairs (positive and negative) was randomly shuffled and then split into training (80 %) and testing (20 %) sets using stratified sampling to preserve the class distribution. From the training set, an additional 20 % was held out as a validation set, again via stratified random sampling. For DDTRN, the validation set was used for early stopping and hyperparameter tuning. The training procedure employed the Adam optimizer with an initial learning rate of 1.5 × 10^−4^, a batch size of 32, and an exponential learning rate scheduler with a decay factor (*γ*) of 0.99 per epoch. The model was trained for up to 300 epochs with early stopping patience of 10 epochs based on validation loss; the model parameters yielding the lowest validation loss were restored. For the baseline methods, the same training/validation/test splits were used. Classifiers supporting early stopping (XGBoost, LGBM, GBDT, and MLP) were provided with the validation set to terminate training when performance ceased to improve. All other classifiers were trained on the training set without validation intervention. The random seed was fixed for all experiments to ensure reproducibility.

### 2.5 Metrics for evaluation

The performance of DDTRN and all baseline methods was assessed using six standard classification metrics computed on the held-out test set (**Figure 1D**). Accuracy was defined as the proportion of correctly classified instances. Precision (positive predictive value) measured the fraction of predicted positive interactions that were true positives, while recall (sensitivity) measured the fraction of actual positive interactions that were correctly identified. The F1 score, the harmonic mean of precision and recall, provided a balanced measure. In addition, two threshold-independent metrics were used: the area under the receiver operating characteristic curve (AUROC) and the area under the precision–recall curve (AUPR). For AUROC and AUPR, the predicted probability of the positive class (regulatory interaction) was used as the decision score. In DDTRN, these probabilities were obtained via the softmax output of the classifier head; in baseline methods, they were obtained either from predict_proba (for classifiers that support it) or from decision_function (for SVM). All metrics were computed using the scikit-learn library.

### 2.6 Interpretability analysis

To interpret the learned features underpinning DDTRN’s predictive performance, we systematically analyzed the two core components of the DD: the Composition Weight Map (CWM, parameter tensor **M**) and the Position Weight Function (PWF, parameter tensor **P**). Both sets of parameters are learned end-to-end during training without additional supervision, and their structures provide direct mechanistic insight into how sequence composition and positional arrangement contribute to regulatory prediction (**Figure 1D**).

For CWM analysis, we focused on the encoding vectors assigned to each possible *r*-mer token (e.g., *rank* = 6 in default settings). For each token, we computed the L2 norm of its corresponding embedding vector as a proxy for its overall influence on the sequence representation. Tokens were then ranked by this norm, and the top-ranking *r*-mers were examined for sequence-level patterns. This analysis aimed to identify which short sequence fragments the model consistently treated as informative for distinguishing regulatory from non-regulatory TF–TG pairs.

For PWF analysis, we examined the learned tensor **P** of shape *m*×*m*×*o*, where each element *P*_*pqh*_ modulates the contribution of a specific cosine basis function with period defined in Eq. (2). To quantify the relative importance of each periodic component, we calculated the absolute value |*P*_*pqh*_| and ranked the corresponding basis terms accordingly. This procedure was performed separately for different rank settings (1 to 6) under both linear and nonlinear token extraction modes. Larger |*P*_*pqh*_| values indicate stronger reliance on particular periodic positional patterns during sequence encoding. By comparing the distribution and magnitude of these values across ranks, we assessed how the model compensates for limited local sequence information with positional weighting.

The interpretability analysis was conducted on the trained DDTRN models using the Dream5_Ecoli dataset. All visualizations, including bar plots for top-ranked *r*-mers and heatmaps for periodic terms, were generated using Matplotlib.

## 3. Results

### 3.1 Performance evaluation

To evaluate the performance of DDTRN in inferring bacterial TRNs, we conducted a comparative analysis against six baseline methods (SVM, RF, GBDT, LGBM, XGBoost, MLP) across the eight datasets. For these machine learning methods, we used *k*-mer frequency vectors as the feature representation for the concatenated sequences. A sliding window with a stride of 1 was used to extract all *k*-mers, count the occurrences of each *k*-mer, and normalize them to frequencies, resulting in a feature vector of size 4^rank^ (where the *k*-mer length corresponds to the ‘rank’ parameter in DDTRN). All methods were evaluated using a consistent evaluation strategy and the same test sets.

As shown in **Table 2**, the predictive performance of DDTRN and six baseline machine learning methods at *rank* = 6 was evaluated across eight benchmark datasets. Overall, DDTRN achieved the best average performance among all methods, with an average AUROC of 0.8690, an average AUPR of 0.8683, and an average F1-score of 0.8053. These results indicate that DDTRN provides the most balanced overall performance across different bacterial TRN datasets. Furthermore, on the *M. tuberculosis* dataset, which contains the largest number of known regulatory interactions among the benchmark datasets, DDTRN consistently outperformed all baseline methods across all metrics. This suggests that DDTRN remains effective in more complex and densely connected regulatory networks. While some baseline methods demonstrate competitiveness on specific datasets, such as SVM on the RegulonDB_Ecoli dataset, and GBDT or LGBM on some other datasets, their strengths are often limited to specific datasets or metrics. In contrast, DDTRN exhibits more stable overall performance across a wide range of datasets, ranking highly in average AUROC, AUPR, and F1 scores. This suggests that the sequence representations learned by DDTRN are more consistently able to capture regulatory information than traditional *k*-mer frequency-based machine learning models.

**Table 2.** Performance of various methods on the eight benchmark datasets.

| Dataset | Metric | DDTRN | SVM | GBDT | LGBM | RF | XGBoost | MLP |
| --- | --- | --- | --- | --- | --- | --- | --- | --- |
| Dream5_Ecoli | AUROC | **0.8213** | 0.8005 | 0.7886 | 0.7861 | 0.7869 | 0.7814 | 0.7620 |
|  | AUPR | **0.8296** | 0.7934 | 0.8085 | 0.8031 | 0.8097 | 0.7997 | 0.7686 |
|  | F1 | **0.7429** | 0.7277 | 0.7141 | 0.7031 | 0.7039 | 0.7177 | 0.6916 |
| RegulonDB_Ec<br>oli | AUROC | 0.8441 | **0.8681** | 0.8494 | 0.8420 | 0.8533 | 0.8486 | 0.8388 |
|  | AUPR | 0.8523 | **0.8726** | 0.8598 | 0.8470 | 0.8626 | 0.8569 | 0.8353 |
|  | F1 | 0.7699 | **0.7892** | 0.7692 | 0.7701 | 0.7579 | 0.7756 | 0.7643 |
| Subtiwiki_Bsubt<br>ilis | AUROC | 0.8299 | **0.8357** | 0.8159 | 0.8169 | 0.8177 | 0.8024 | 0.7868 |
|  | AUPR | 0.8309 | **0.8465** | 0.8259 | 0.8180 | 0.8230 | 0.8063 | 0.8004 |
|  | F1 | **0.7578** | 0.7454 | 0.7298 | 0.7506 | 0.7201 | 0.7325 | 0.7138 |
| *S. enterica* | AUROC | **0.8703** | 0.8450 | 0.8568 | 0.8435 | 0.8597 | 0.8453 | 0.8060 |
|  | AUPR | **0.8881** | 0.8702 | 0.8800 | 0.8657 | 0.8831 | 0.8729 | 0.8314 |
|  | F1 | **0.8063** | 0.7953 | 0.7674 | 0.7734 | 0.7680 | 0.7616 | 0.7535 |
| *C. glutamicum* | AUROC | 0.9218 | 0.9224 | 0.9216 | **0.9245** | 0.9211 | 0.9128 | 0.9035 |
|  | AUPR | **0.9366** | 0.9258 | 0.9304 | 0.9334 | 0.9329 | 0.9289 | 0.9169 |
|  | F1 | 0.8490 | **0.8546** | 0.8472 | 0.8464 | 0.8449 | 0.8402 | 0.8305 |
| *M. tuberculosis* | AUROC | **0.8263** | 0.8227 | 0.8145 | 0.7873 | 0.8054 | 0.8005 | 0.8090 |
|  | AUPR | **0.7918** | 0.7887 | 0.7895 | 0.7646 | 0.7716 | 0.7766 | 0.7818 |
|  | F1 | **0.7679** | 0.7665 | 0.7420 | 0.7185 | 0.7363 | 0.7335 | 0.7365 |
| *P. aeruginosa* | AUROC | 0.8885 | 0.8823 | **0.8906** | 0.8861 | 0.8827 | 0.8800 | 0.8617 |
|  | AUPR | 0.8756 | 0.8645 | **0.8886** | 0.8744 | 0.8757 | 0.8624 | 0.8305 |
|  | F1 | **0.8327** | 0.8243 | 0.8211 | 0.8229 | 0.8167 | 0.8134 | 0.8078 |
| *S. coelicolor* | AUROC | 0.9497 | 0.9372 | 0.9520 | **0.9521** | 0.9502 | 0.9508 | 0.9410 |
|  | AUPR | 0.9417 | 0.9209 | **0.9495** | 0.9407 | 0.9343 | 0.9437 | 0.9218 |
|  | F1 | 0.9162 | 0.8979 | 0.9127 | **0.9178** | 0.9157 | 0.9120 | 0.8937 |
| **Average** | AUROC | **0.8690** | 0.8642 | 0.8612 | 0.8548 | 0.8596 | 0.8527 | 0.8386 |
|  | AUPR | **0.8683** | 0.8603 | 0.8665 | 0.8559 | 0.8616 | 0.8559 | 0.8358 |
|  | F1 | **0.8053** | 0.8001 | 0.7879 | 0.7878 | 0.7829 | 0.7858 | 0.7740 |

Furthermore, the results in supplementary **Figure S3** show that DDTRN has a more significant advantage at lower ranks (*rank* = 1, 2, 3). This trend can be explained by differences in sequence representation strategies. Traditional machine learning methods use *k*-mer frequency vectors to represent tandem TF-TG sequences; these vectors primarily encode compositional information while largely ignoring positional arrangement. Therefore, these methods have limited ability to capture long-range sequence structure at lower ranks. In contrast, the DD representation used in DDTRN integrates CWM and PWF components, enabling the model to simultaneously encode nucleotide composition and positional information. Learnable PWF components assign different weights to different sequence positions, thus compensating for the limited structural information contained in short *k*-mers. As the rank increases, the *k*-mer fragments themselves contain more permutation information, allowing traditional *k*-mer frequency-based methods to gradually improve. Therefore, the performance gap between DDTRN and baseline methods gradually narrows at higher ranks. Subsequent PWF parameter analysis also supports this explanation: at low ranks, the |*P*_*pqh*_| values of the first few terms are significantly larger, indicating a greater reliance on positional weights to enhance sequence encoding; as the rank increases, the |*P*_*pqh*_| values generally stabilize, and the contribution of PWF gradually weakens.

### 3.2 Parameter analysis

The performance of the DDTRN model is influenced by several parameters, including descriptor rank (*rank*), vector embedding dimension (*vec_dim*), and the number of basis functions used for expanding the position weight function (*num_basis*). To assess the model’s effectiveness in predicting TRNs, we compared the trends of AUROC, AUPR, and F1 across the eight benchmark datasets, varying one parameter while keeping the other two constant to adhere to the principle of univariate analysis. We conducted fine-tuning by adjusting parameter values to thoroughly analyze their impact on TRN prediction.

#### 3.2.1 Sensitivity analysis of rank

In DDTRN, rank determines the length of the sequence fragments (tokens) extracted from tandem TF-TG gene sequences. When *rank* = *k*, the model uses *k*-mer level tokens to represent sequences and applies weights to these tokens based on their position before aggregating them into a description vector. Therefore, the value of rank directly affects the model’s ability to capture local sequence patterns related to regulatory relationships.

To investigate the impact of this parameter, we evaluated DDTRN with rank values of 1, 3, 6, 9, and 11 on eight benchmark datasets. As shown in **Figure 2**, performance generally improves significantly as rank increases from 1 to 6. This trend was observed on AUROC, AUPR, and F1 metrics across all datasets, indicating that short tokens with low rank values provide insufficient local sequence information, while *k*-mers of moderate length are more effective in capturing regulatory sequence patterns in tandem TF-TG gene sequences. When rank is further increased from 6 to 9 or 11, the performance improvement decreases significantly and tends to stabilize. This suggests that beyond a certain limit, increasing token length does not consistently introduce proportionally effective information. Instead, longer *k*-mers may increase feature sparsity, generating more rare or noisy token combinations and increasing computational complexity.

**Figure 2.**
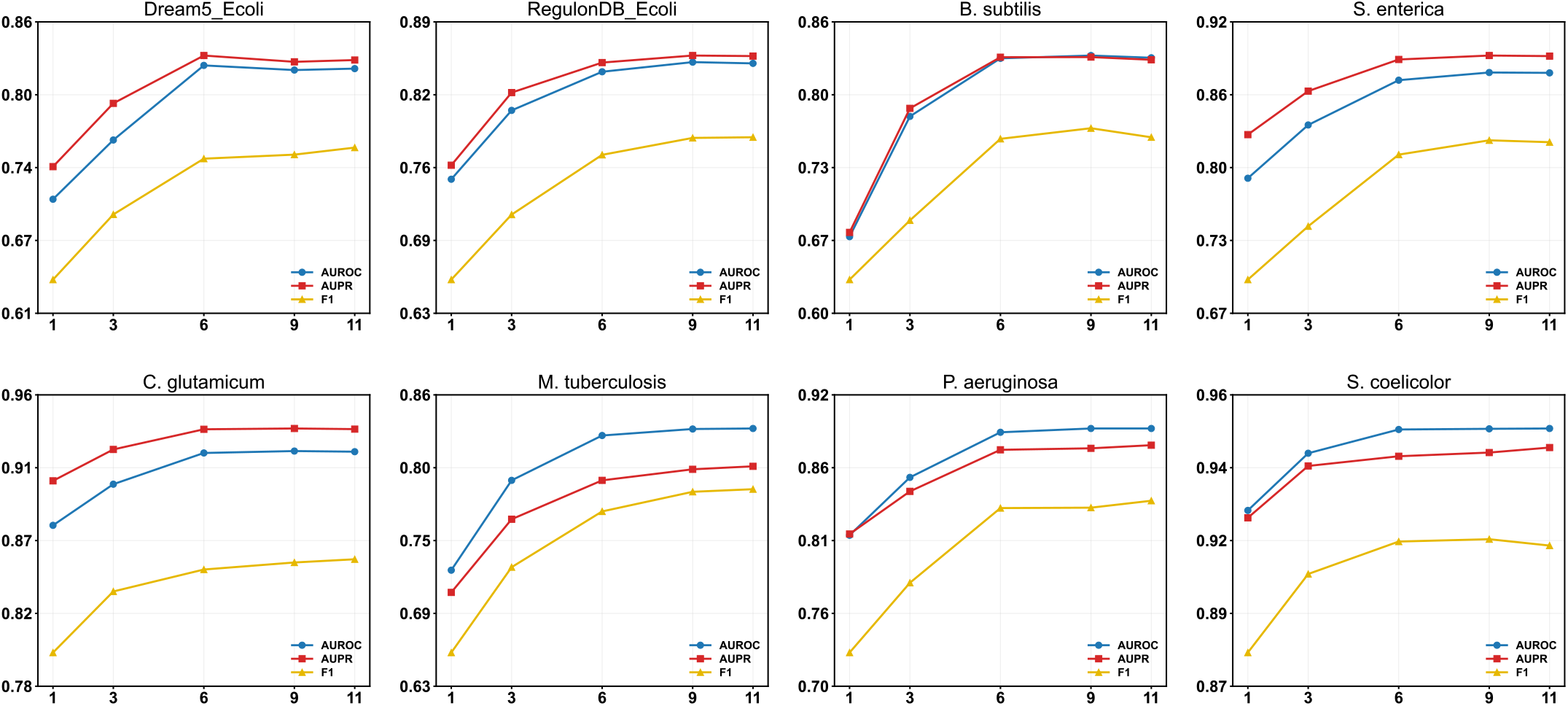
Sensitivity analysis for parameter *rank*. Performance generally increases with rank and *rank* = 6 is a good choice and adopted to show the prediction performance.

Considering both prediction performance and computational efficiency, we choose *rank* = 6 as the default setting for DDTRN. This value captures sufficient local sequence information while avoiding increased computational complexity due to excessively high rank values. Therefore, *rank* = 6 represents the optimal choice for model accuracy, stability, and computational cost in subsequent experiments.

#### 3.2.2 Sensitivity analysis of vec_dim

The vec_dim parameter determines the dimensionality of the encoding vector representation in DDTRN and affects the expressive capacity of the sequence descriptor. We evaluated vec_dim values of 2, 4, 6, 8, 10, and 12 across eight datasets. As shown in **Figure S1**, performance generally improved when vec_dim increased from 2 to 4 or 6, indicating that very low-dimensional embeddings are insufficient to capture regulatory sequence patterns. After vec_dim reached 6–8, AUROC, AUPR, and F1 tended to stabilize in most datasets, and further increasing the dimension to 10 or 12 only brings a slight improvement. Therefore, we selected *vec_dim* = 10 as the default setting, as it provides stable near-optimal performance while maintaining sufficient representational capacity and acceptable computational complexity.

#### 3.2.3 Sensitivity analysis of num_basis

The num_basis parameter controls the number of basis functions used to construct the positional weight function (PWF), thus affecting the granularity of positional encoding in DDTRN. We evaluated num_basis values of 2, 4, 6, 8, 10, and 12 on eight datasets. As shown in **Figure S2**, the AUROC, AUPR, and F1 scores change little with the num_basis value, indicating that DDTRN is relatively insensitive to this parameter and maintains stable performance over a wide range of basis function numbers. While no sustained monotonic improvement was observed, the F1 score improved slightly on some datasets when num_basis was increased to around 10, such as Dream5_Ecoli, Subtiwiki_Bsubtilis, *S. enterica*, and *P. aeruginosa*. Further increasing num_basis to 12 did not yield significant additional improvements and even showed slight fluctuations on some datasets. Therefore, we chose *num_basis* = 10 as the default setting for this study.

#### 3.2.4 Robustness of DDTRN

To evaluate the impact of training data volume on model performance, we randomly sampled 10%, 20%, 30%, 40%, 50%, and 60% of the training data from eight datasets, while maintaining the independence of the validation and test sets. Each setting was repeated using a different random seed, and box plots were used to summarize AUROC and AUPR values. As shown in **Figure 3**, DDTRN maintained stable performance on most datasets as the training data size varied. On some datasets, even at lower training ratios, the model achieved relatively high AUROC and AUPR values, and the results of repeated runs were relatively concentrated. For Dream5_Ecoli, Subtiwiki_Bsubtilis, and *S. enterica* datasets, performance significantly improved when the training ratio increased from 10% to 20% or 30%, then gradually stabilized. These results demonstrate good robustness and performance even on a small number of sampled training datasets.

**Figure 3.**
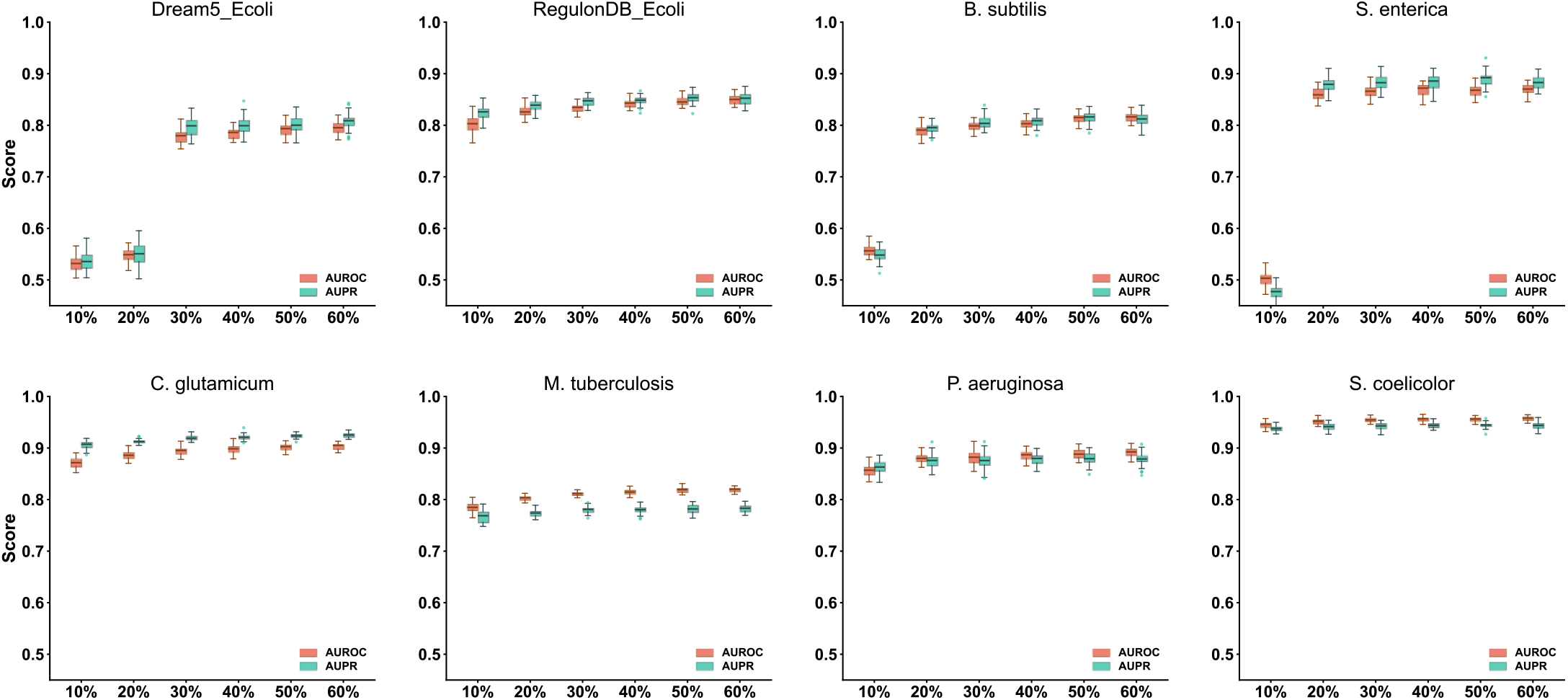
Robustness analysis for data size. Generally, DDTRN is not sensitive to data size variation, and on most datasets, 20% to 30% sample size is enough for good prediction.

### 3.3 Interpretability analysis

### 3.3.1 Interpretation of CWM

In DDTRN, M corresponds to the learnable parameters of the Composition Weight Map (CWM), which assigns a vector representation to each token of length rank. When *rank* = 6, there are 4^6^=4096 possible 6-mers, and each 6-mer is associated with a learnable embedding vector. To estimate the relative contribution of different 6-mers, we calculated the L2 norm of each embedding vector and ranked all 6-mers accordingly. As shown in **Figure 4**, the top 20 6-mers with the largest M values were identified under the nonlinear mode at *rank* = 6. The CWM parameter M mainly captures compositional patterns that contribute strongly to sequence representation. The top-ranked 6-mers, such as GCGACG, GTAGTT, AGTATG, and CGCGTG, exhibited relatively large embedding norms, indicating that these fragments had stronger influence on the learned sequence representation.

**Figure 4.**
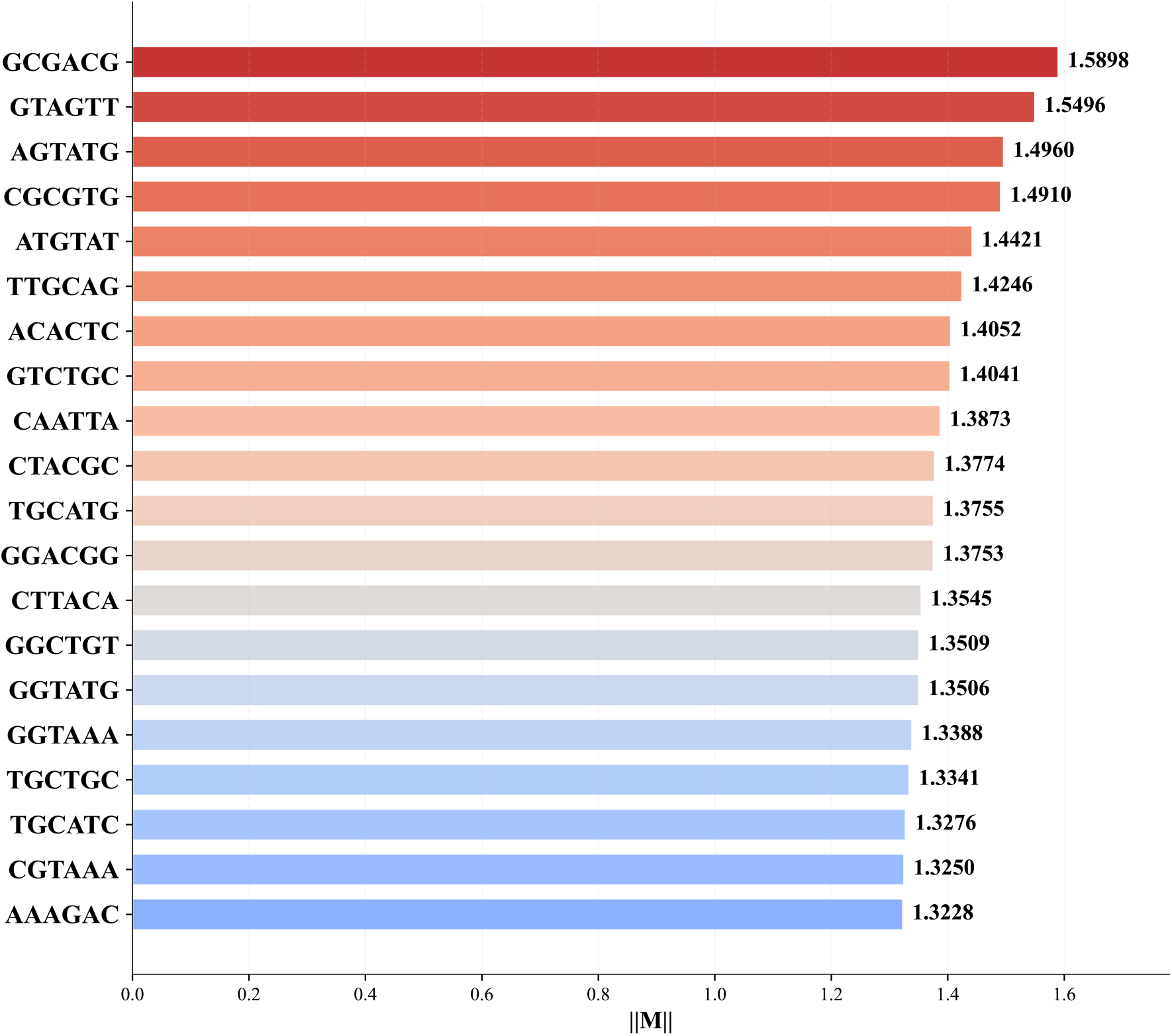
The top 20 most influential 6-mer tokens learned by CWM. The bar chart represents the length (norm) of each 6-mer embedding vector.

To characterize these high-norm 6-mers, we translated the top-ranked 6-mer tokens into amino-acid residues and examined whether specific residue categories were enriched. As shown in **Figure 5**, using the *E. coli* genomic amino-acid background calculated from coding sequences, the combined frequency of Cys and Met was 3.97% (53,680/1,353,650). In contrast, the translated residues of the top20 6-mers contained 10 Cys/Met residues out of 40 total residues, whereas only 1.59 residues would be expected based on the *E. coli* genome background. This corresponds to a 6.3-fold enrichment, with a significant enrichment signal after multiple-testing correction (one-sided binomial test *p* = 3.00 × 10^−6^, *FDR* = 1.40 × 10^−5^). Similar significant enrichment was also observed in larger top-ranked sets (e.g., top50, top100), suggesting that sulfur-containing amino-acid-associated codon patterns are preferentially represented among the highest-weighted CWM tokens. The biophysical/biochemical mechanisms underlying this enrichment require further investigation to be clarified.

**Figure 5.**
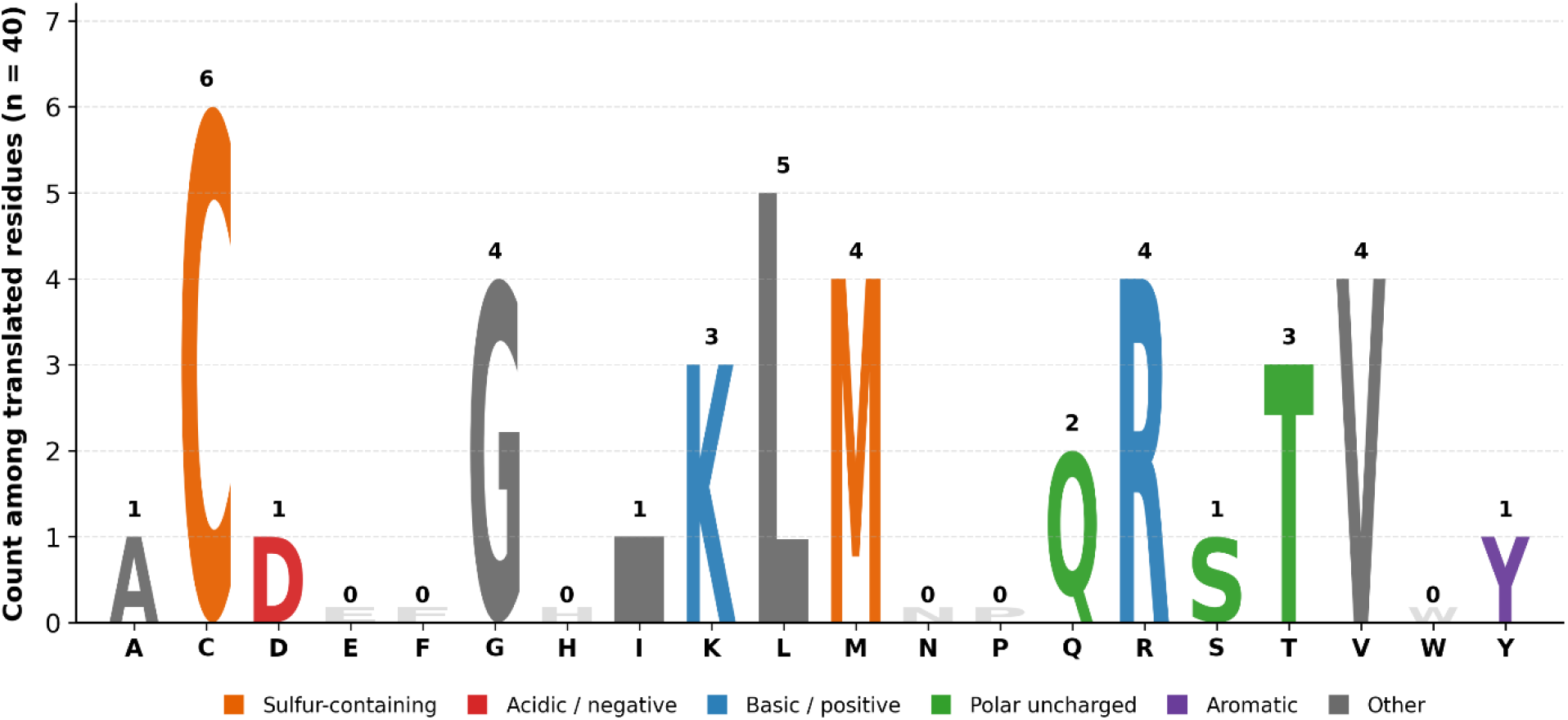
Amino-acid composition of translated top 20 6-mers. Letter height represents the count of each amino acid among 40 translated residues. Cys and Met are highlighted in orange, showing significant enrichment of sulfur-containing residues in the top-ranked CWM tokens.

#### 3.3.2 Interpretation of PWF

In DDTRN, **P** represents the learnable parameter tensor associated with the Position Weight Functions (PWFs), which encodes position-dependent information in sequence representation. To investigate how PWF contributes under different rank settings, we analyzed the learned |*P*_*pqh*_| values on the Dream5_Ecoli dataset from rank 1 to rank 6, using it to measure the relative contribution strength of each periodic basis term. Different from the main benchmark experiments, the mode was set to “nonlinear” in this analysis. In nonlinear mode, tokens of length rank are extracted without redundant sliding windows, so |*P*_*pqh*_| reflects the contribution of positional information at the token level rather than at each single-base position. Each coefficient *P*_*pqh*_ corresponds to a cosine basis term with a period defined in Eq. (2). Therefore, a larger |*P*_*pqh*_| value indicates that the corresponding periodic component contributes more strongly to sequence encoding. **Supplementary File 1** provides statistics for the |*P*_*pqh*_| values of periodic terms 2-1001 across different ranks.

**Figure 6** displays the top 20 periodic terms ranked by |*P*_*pqh*_| magnitude for ranks 1 through 6. It can be seen that at lower ranks (*rank* =1, 2, 3), the |*P*_*pqh*_| values for the top 20 terms are significantly higher than those at higher ranks (*rank* = 4, 5, 6). This suggests that when the local contextual information carried by each low-rank fragment is limited, the model relies more heavily on high-magnitude periodic terms to encode the sequence. As rank increases, the overall magnitude of |*P*_*pqh*_| tends to stabilize because higher-rank fragments inherently contain richer sequence information, reducing the model’s reliance on specific periodic contributions for enhancement. This also corroborates that at higher ranks, the performance gain from introducing positional information via PWF in the DD model is relatively limited, thereby narrowing the advantage over ML methods based solely on *k*-mer frequency features. Additionally, significant differences in |*P*_*pqh*_| values among the top 20 periodic terms across different ranks are observable, indicating that the model captures distinct sequence structural features at different rank granularities. For completeness of the analysis, **Figure S4** and **Supplementary File 2** also provide the |*P*_*pqh*_| value analysis for the linear mode.

### 3.4 Case demonstration

To further compare the prediction characteristics of DDTRN with those of conventional machine learning models, we selected the three TF genes with the largest numbers of known regulatory relationships in the RegulonDB_Ecoli dataset, namely *crp, nac*, and *lrp*, for case study analysis. For each TF, we first compared the top 100 predicted positive targets by DDTRN, SVM, and GBDT. As shown in **Figure 7A**, the limited intersections among the top-ranked predictions of the three methods indicate that they prioritize candidate regulatory relationships differently. In particular, the intersections between DDTRN and the two conventional machine learning methods were generally smaller than the intersection between SVM and GBDT, suggesting that SVM and GBDT exhibit relatively similar ranking preferences in this task. These results indicate that DDTRN exhibits complementary prioritization of high-ranking candidate regulatory relationships relative to conventional *k*-mer-based ML models.

**Figure 6.**
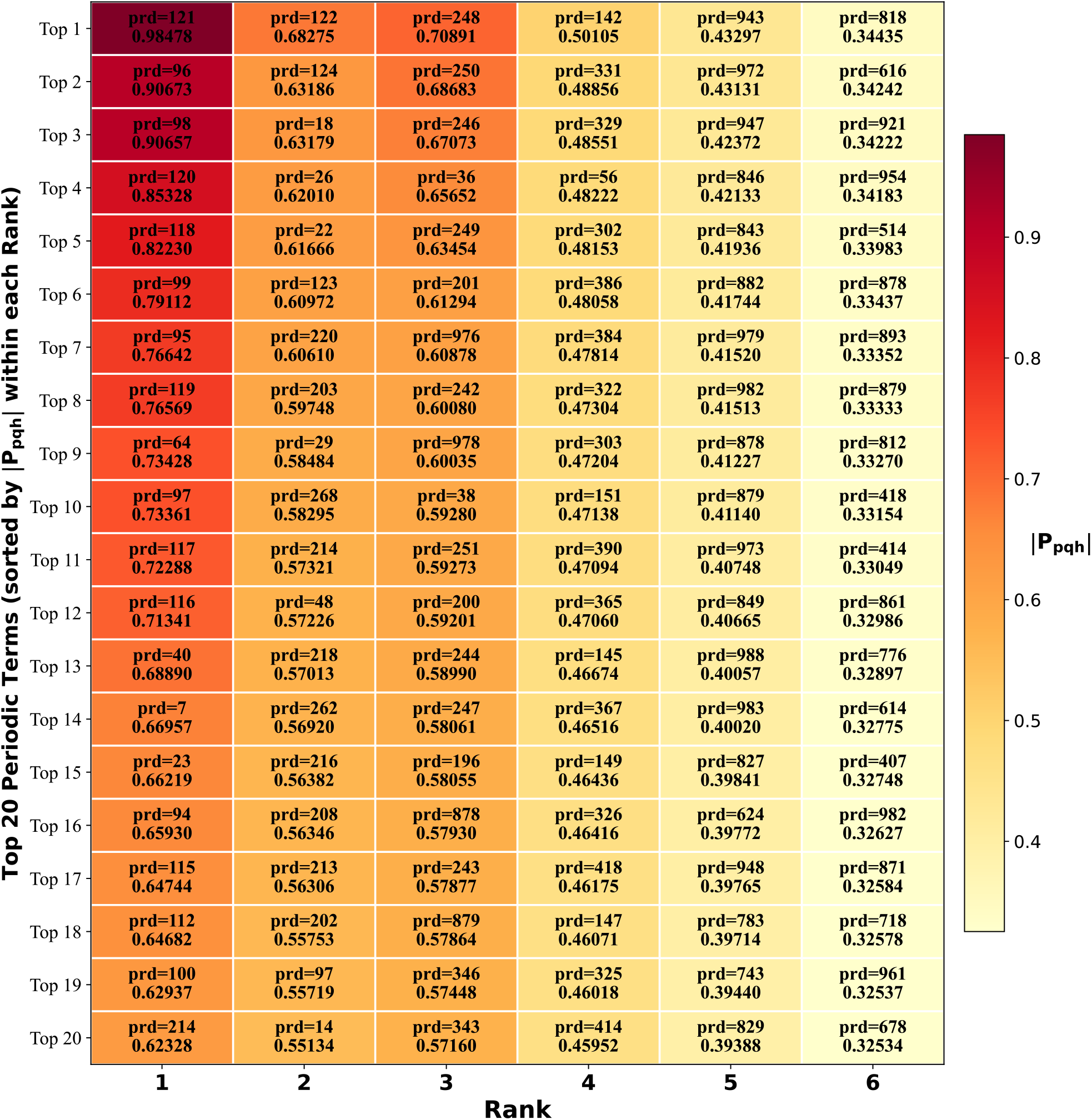
Distribution of |*P*_*pqh*_| values for periodic terms in the PWF of DD across different rank settings (nonlinear mode). Heatmap shows the top 20 periodic terms sorted by the absolute value of the elements in tensor **P** (namely |*P*_*pqh*_|) across ranks 1 to 6. “prd” indicates the corresponding periodic terms defined in Eq. (2). Larger |*P*_*pqh*_| value indicates greater contribution from the corresponding periodicity in the sequence pattern differentiating Regulatory and Non-regulatory interaction.

**Figure 7.**
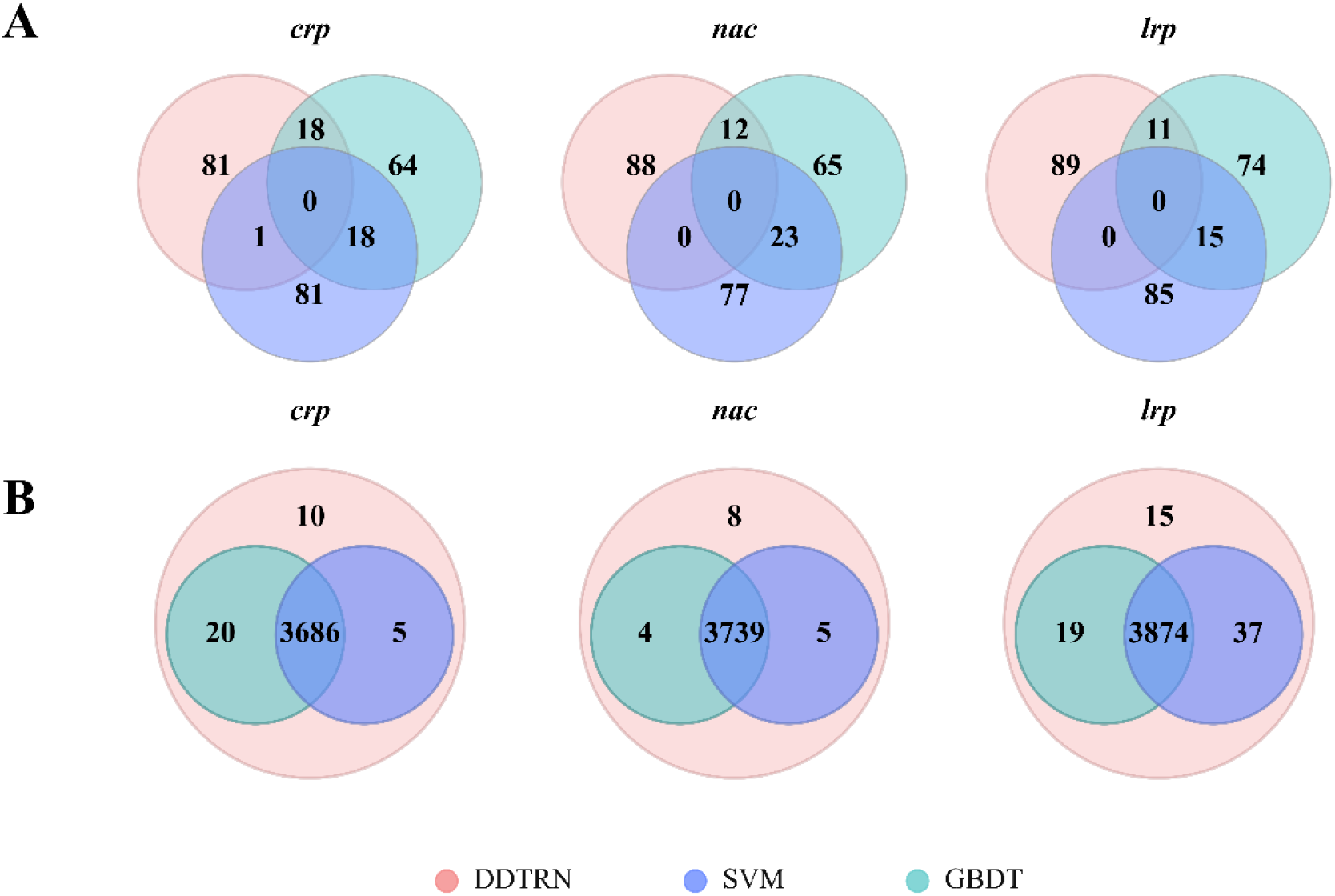
Comparison of top-ranked and overall predicted regulatory targets for TF genes *crp, nac*, and *lrp*. Venn diagrams comparing predicted positive targets by DDTRN, SVM, and GBDT for the three TF genes with the largest numbers of known regulatory relationships in the RegulonDB_Ecoli dataset: *crp, nac*, and *lrp*.**(A)** Comparison of the top 100 (ranked by model score) predicted targets for each TF gene. **(B)** Comparison of all predicted positive regulatory relationships for each TF. The numbers indicate the counts of candidate target genes in each model-specific or shared region.

When all predicted positive regulatory relationships were considered, a large proportion of candidate interactions were shared by the three methods, indicating overall consistency among their prediction results (**Figure 7B**). Meanwhile, DDTRN also identified model-specific candidate regulatory relationships beyond those predicted by SVM and GBDT. These additional predictions may reflect the ability of DDTRN to extract more comprehensive sequence information by integrating both compositional and positional features, thereby capturing regulatory signals that cannot be fully represented by *k*-mer frequency statistics alone.

### 3.5 Overall prediction

Previous network inference studies have shown that no single inference method performs optimally across all datasets, whereas integrating predictions from multiple methods can improve robustness across diverse datasets [20]. Inspired by this, we retained TF–TG pairs predicted as positive by DDTRN, SVM, and GBDT as higher-confidence candidate regulatory relationships. For *E. coli*, RegulonDB_Ecoli instead of Dream5_Ecoli was selected as the representative dataset, because it contains a larger number of known regulatory relationships, making it more suitable for candidate regulatory relationship mining and consensus screening.

As shown in **Table 3**, the number of potential regulatory relationships predicted by each model varied across datasets, reflecting differences in feature representation and classification strategies among DDTRN, SVM, and GBDT. To obtain more reliable candidate interactions, we focused on the “All3 Consensus” results, namely TF–TG pairs simultaneously predicted as positive by all three models. The consensus predictions ranged from 4,279 pairs in *C. glutamicum* to 89,411 pairs in *B. subtilis*, with 87,624 consensus pairs identified in RegulonDB_Ecoli and 84,420 in *M. tuberculosis*. These jointly predicted relationships are less likely to be driven by the bias of a single model and therefore represent higher-confidence candidates for subsequent experimental validation, TRN annotation, and functional analysis. Complete prediction results are provided in **Supplementary Files 3–9**.

**Table 3.**
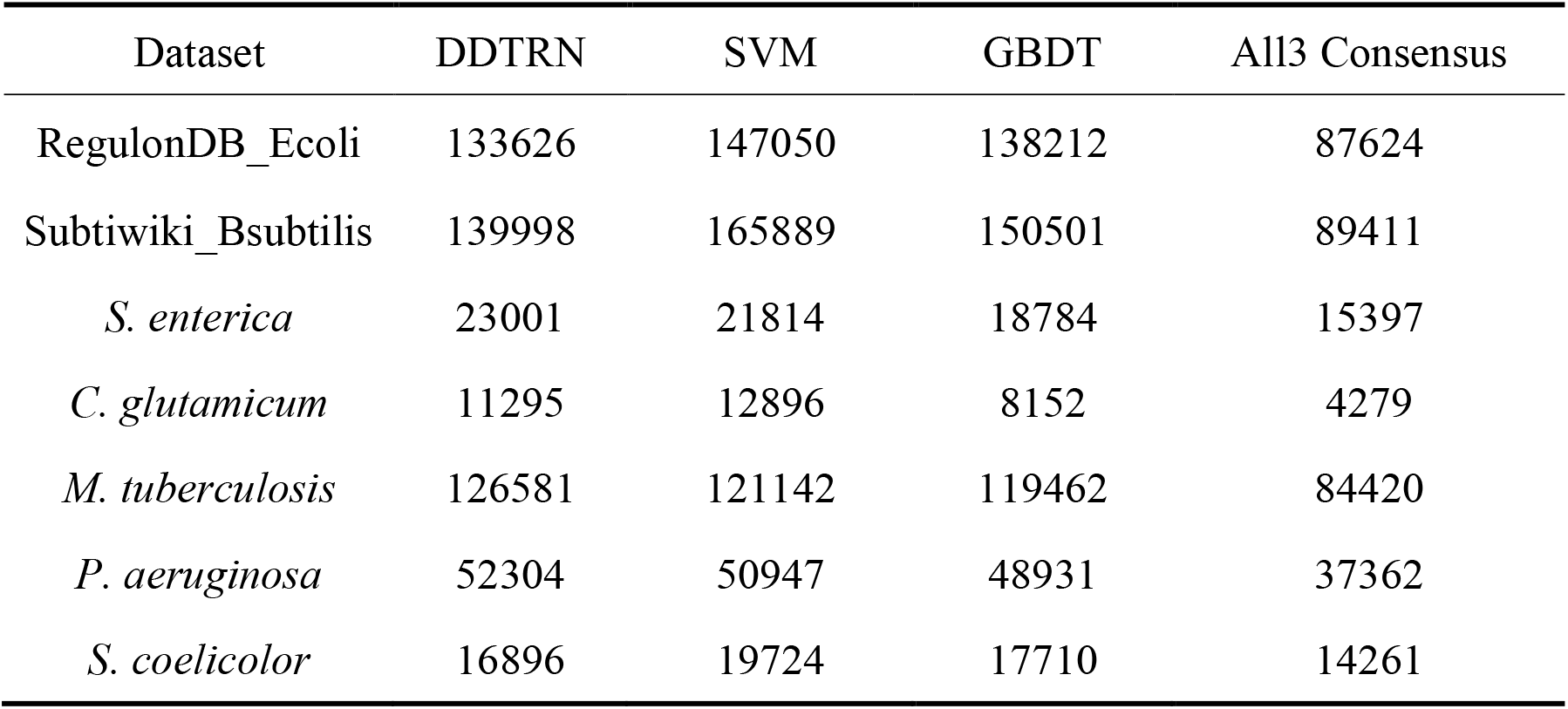
The number of regulatory relationships predicted by the three models across seven datasets.

## 4. Discussion

In this study, we proposed DDTRN, a sequence-driven framework for predicting bacterial transcriptional regulatory networks based on the DD model. Existing TRN inference methods have commonly relied on transcriptomic profiles, graph-based representations, or integrated omics features. For example, previous network inference studies have shown that combining predictions from multiple methods can improve robustness, and recent bacterial TRN models have incorporated expression data and genomic information for supervised regulatory prediction [12, 14]. However, transcriptome-based inference can still be affected by data quality, noise, batch effects, and condition specificity, and benchmark studies have shown that the inference performance of gene regulatory network is sensitive to data type and evaluation settings [32]. Therefore, DDTRN provides a complementary strategy based on gene sequences, especially suitable for bacterial species with limited expression data.

Across eight bacterial benchmark datasets, DDTRN achieved the best average performance among all compared methods, with an average AUROC of 0.8690, AUPR of 0.8683, and F1-score of 0.8053. Compared with conventional machine learning methods based on *k*-mer frequency features, DDTRN showed more stable overall performance across datasets. This result supports the idea that genomic sequences contain regulatory-relevant information. Previous sequence-based models, such as DeepBind, DeepSEA, DanQ, and Enformer, have demonstrated that DNA sequence can encode information related to binding specificity, regulatory sequence code, motif grammar, and gene expression patterns [15-18]. However, compared with many deep learning models whose internal representations are often difficult to interpret, DDTRN retains a more explicit and interpretable representation through the DD architecture. Specifically, the CWM captures compositional information of sequence tokens, whereas the PWF introduces position-dependent or arrangement-related information, consistent with the theoretical basis of the DD method [25].

The parameter sensitivity analyses showed that DDTRN maintained stable performance under different settings of rank, vec_dim, and num_basis. In particular, DDTRN showed a more obvious advantage at lower ranks, where short *k*-mers contain limited context information. This suggests that PWF can compensate for insufficient context information by introducing position-dependent weighting. As rank increases, *k*-mer tokens themselves contain richer sequence patterns, reducing the relative contribution of PWF and narrowing the performance gap between DDTRN and conventional *k*-mer frequency-based methods. The robustness analysis further showed that DDTRN can maintain reliable predictive performance when the amount of training data is reduced, suggesting its potential utility in limited-sample scenarios.

In addition to predictive performance, DDTRN provides interpretable model parameters. The PWF analysis showed that positional periodic components contributed differently across rank settings, with stronger contributions at lower ranks and more stable values at higher ranks. This supports the view that PWF provides additional positional information when the token length is short. The CWM analysis further identified high-norm 6-mer tokens that contributed strongly to sequence representation. Translation-based composition analysis further showed that the top20 high-norm 6-mers were significantly enriched in sulfur-containing amino-acid-associated codon patterns, suggesting a non-random compositional preference among the strongest CWM tokens. The preference in amino acid composition may be associated with certain structural features of TFs [33], and the underlying molecular mechanisms require further investigation.

We also applied DDTRN together with SVM and GBDT to predict candidate TF–TG relationships across seven datasets and identified consensus regulatory relationships supported by all three models. Because DDTRN, SVM, and GBDT use different sequence representations and classification strategies, their shared positive predictions are more likely to represent cross-model regulatory signals rather than model-specific biases. In addition, curated bacterial regulatory resources such as RegulonDB, RegPrecise, DBTBS, and CoryneRegNet show that bacterial TRN annotation is a continuous process that combines literature curation, experimental evidence, and comparative genomics [34-37]. Therefore, the consensus TF–TG pairs identified in this study may serve as prioritized candidates for future experimental validation and bacterial TRN annotation.

Despite these advantages, several limitations of DDTRN should be acknowledged. First, bacterial transcriptional regulation is influenced by a variety of factors, including TF activity, environmental signals, cofactors, promoter architecture, nucleoid-associated proteins, and combinatorial regulation [38]. Therefore, sequence information alone cannot fully reflect the dynamic regulatory state of a cell. Second, current bacterial TRN annotations remain incomplete even in well-curated resources, and negative samples generated from currently unannotated TF–TG pairs may include undiscovered regulatory interactions. This limitation may constrain supervised learning performance. Third, the current DDTRN framework uses concatenated gene sequences rather than experimentally defined promoter regions or TF-binding regions. Future work should incorporate more precise regulatory regions, such as promoter sequences, upstream regions around transcription start sites, or experimentally supported TFBSs. In the future, DDTRN can be extended from a sequence-driven framework to a multimodal regulatory prediction framework, integrating multi-omics and transcriptional regulatory data, which is expected to further improve the accuracy and biological relevance of bacterial TRN reconstruction.

Overall, DDTRN provides a scalable, interpretable, and data-efficient sequence-based framework for bacterial TRN prediction. It complements existing expression-based and graph-based approaches by enabling regulatory relationship prediction in species where large-scale omics data (such as transcriptome and proteome) are limited. With further integration of experimentally defined regulatory regions and multimodal regulatory evidence, DDTRN has the potential to become a more powerful tool for bacterial TRN annotation, candidate regulatory interaction discovery, and functional genomics studies.

## Data availability

The codes and datasets used in this study can be found on GitHub (https://github.com/mbglab/DualDescriptor/tree/main/Apps/DDTRN).

## References

1. Cramer P. Organization and regulation of gene transcription, Nature 2019;573:45–54.

2. Ho HI, Fang JR, Cheung J et al. Programmable CRISPR-Cas transcriptional activation in bacteria, Mol Syst Biol 2020;16:e9427.

3. Schlitt T, Brazma A. Modelling gene networks at different organisational levels, FEBS Lett 2005;579:1859–1866.

4. Vázquez A, Dobrin R, Sergi D et al. The topological relationship between the large-scale attributes and local interaction patterns of complex networks, Proc Natl Acad Sci U S A 2004;101:17940–17945.

5. van Noort V, Snel B, Huynen MA. The yeast coexpression network has a small-world, scale-free architecture and can be explained by a simple model, EMBO Rep 2004;5:280–284.

6. Moore MJ. From Birth to Death: The Complex Lives of Eukaryotic mRNAs, Science 2005;309:1514–1518.

7. Zhao J, Sun X, Mao Z et al. Independent component analysis of Corynebacterium glutamicum transcriptomes reveals its transcriptional regulatory network, Microbiological Research 2023;276:127485.

8. The ENCODE (ENCyclopedia Of DNA Elements) Project, Science 2004;306:636–640.

9. Dong J, Li J, Wang F. Deep Learning in Gene Regulatory Network Inference: A Survey, IEEE/ACM Transactions on Computational Biology and Bioinformatics 2024;21:2089–2101.

10. Park PJ. ChIP-seq: advantages and challenges of a maturing technology, Nat Rev Genet 2009;10:669–680.

11. Bartlett A, O’Malley RC, Huang SC et al. Mapping genome-wide transcription-factor binding sites using DAP-seq, Nat Protoc 2017;12:1659–1672.

12. Razaghi-Moghadam Z, Nikoloski Z. Supervised learning of gene-regulatory networks based on graph distance profiles of transcriptomics data, NPJ Syst Biol Appl 2020;6:21.

13. Wang J, Ma A, Ma Q et al. Inductive inference of gene regulatory network using supervised and semi-supervised graph neural networks, Comput Struct Biotechnol J 2020;18:3335–3343.

14. Gu W-C, Ma B-G. PGBTR: a powerful and general method for inferring bacterial transcriptional regulatory networks, BMC Genomics 2025;26:712.

15. Alipanahi B, Delong A, Weirauch MT et al. Predicting the sequence specificities of DNA- and RNA-binding proteins by deep learning, Nature Biotechnology 2015;33:831–838.

16. Zhou J, Troyanskaya OG. Predicting effects of noncoding variants with deep learning–based sequence model, Nature Methods 2015;12:931–934.

17. Quang D, Xie X. DanQ: a hybrid convolutional and recurrent deep neural network for quantifying the function of DNA sequences, Nucleic Acids Research 2016;44:e107–e107.

18. Avsec Ž, Agarwal V, Visentin D et al. Effective gene expression prediction from sequence by integrating long-range interactions, Nature Methods 2021;18:1196–1203.

19. Ma B-G. Analytic Number Theory Model for character sequence and its application in bioinformatics. [Master Thesis] Tian Jing University, 2003. (In Chinese)

20. Marbach D, Costello JC, Küffner R et al. Wisdom of crowds for robust gene network inference, Nat Methods 2012;9:796–804.

21. Catoiu EA, Krishnan J, Li G et al. iModulonDB 2.0: dynamic tools to facilitate knowledge-mining and user-enabled analyses of curated transcriptomic datasets, Nucleic Acids Research 2025;53:D99–D106.

22. Escorcia-Rodríguez JM, Tauch A, Freyre-González JA. Abasy Atlas v2.2: The most comprehensive and up-to-date inventory of meta-curated, historical, bacterial regulatory networks, their completeness and system-level characterization, Comput Struct Biotechnol J 2020;18:1228–1237.

23. O’Leary NA, Cox E, Holmes JB et al. Exploring and retrieving sequence and metadata for species across the tree of life with NCBI Datasets, Scientific Data 2024;11:732.

24. Ma B-G. Dual Descriptor for Sequence Analysis: A Revisit 2025. https://github.com/mbglab/DualDescriptor/blob/main/Refs/DD_Methodology_2025.pdf

25. Ma B-G. Building Block and Building Rule: Dual Descriptor Method for Biological Sequence Analysis, Nature Precedings 2008. https://www.nature.com/articles/npre.2008.2223.1

26. Cortes C, Vapnik V. Support-vector networks, Machine Learning 1995;20:273–297.

27. Breiman L. Random Forests, Machine Learning 2001;45:5–32.

28. Friedman JH. Greedy function approximation: A gradient boosting machine, The Annals of Statistics 2001;29:1189–1232, 1144.

29. Ke G, Meng Q, Finley T et al. LightGBM: A Highly Efficient Gradient Boosting Decision Tree. In: Neural Information Processing Systems. 2017.

30. Chen T, Guestrin C. XGBoost: A Scalable Tree Boosting System, ACM 2016.

31. Rumelhart DE, Hinton GE, Williams RJ. Learning representations by back-propagating errors, Nature 1986;323:533–536.

32. Pratapa A, Jalihal AP, Law JN et al. Benchmarking algorithms for gene regulatory network inference from single-cell transcriptomic data, Nature Methods 2020;17:147–154.

33. Fernandez-Lopez R, Ruiz R, del Campo I et al. Structural basis of direct and inverted DNA sequence repeat recognition by helix–turn–helix transcription factors, Nucleic Acids Research 2022;50:11938–11947.

34. Salgado H, Gama-Castro S, Lara P et al. RegulonDB v12.0: a comprehensive resource of transcriptional regulation in E. coli K-12, Nucleic Acids Research 2024;52:D255–D264.

35. Novichkov PS, Laikova ON, Novichkova ES et al. RegPrecise: a database of curated genomic inferences of transcriptional regulatory interactions in prokaryotes, Nucleic Acids Research 2010;38:D111–D118.

36. Sierro N, Makita Y, de Hoon M et al. DBTBS: a database of transcriptional regulation in Bacillus subtilis containing upstream intergenic conservation information, Nucleic Acids Research 2008;36:D93–D96.

37. Parise MTD, Parise D, Kato RB et al. CoryneRegNet 7, the reference database and analysis platform for corynebacterial gene regulatory networks, Scientific Data 2020;7:142.

38. Balleza E, López-Bojorquez LN, Martínez-Antonio A et al. Regulation by transcription factors in bacteria: beyond description, FEMS Microbiol Rev 2009;33:133–151.

