## Supplementary Data for "DDTRN: Predicting Bacterial Transcriptional Regulatory Networks Based on Gene Sequences using Dual Descriptor"

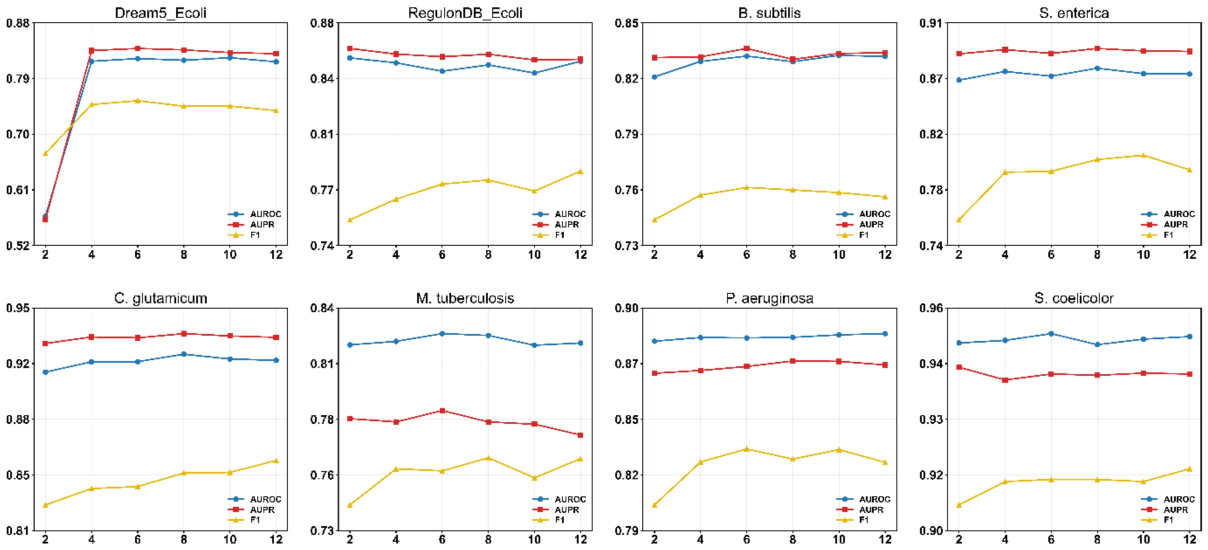


**Figure S1. Sensitivity analysis of the vec_dim parameter.** Performance generally increases with vec_dim and *vec_dim* = 10 is adopted to show the prediction performance.

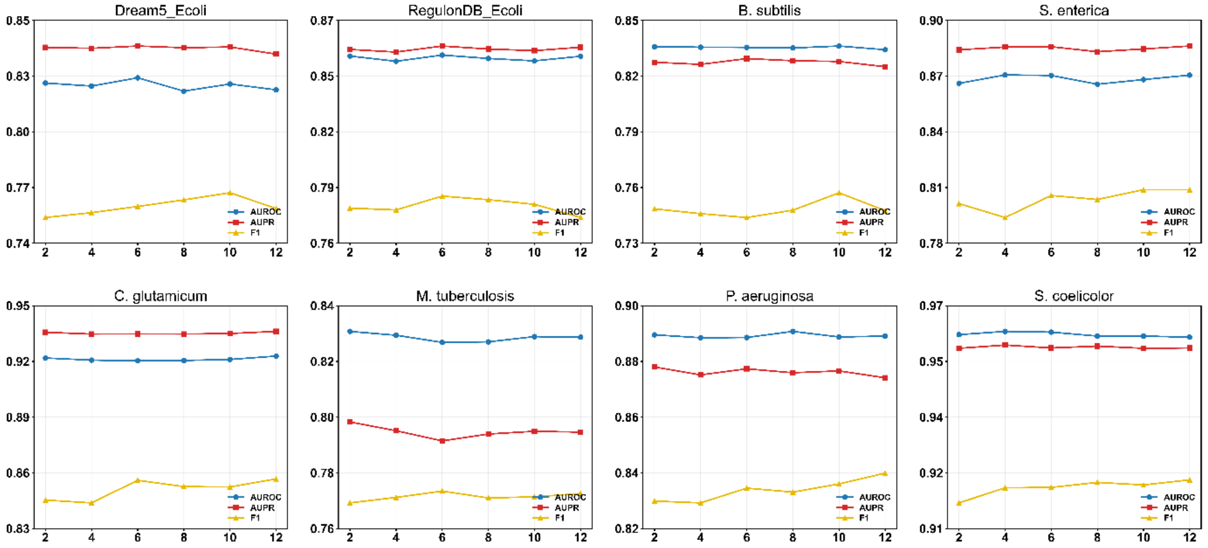


**Figure S2. Sensitivity analysis of the num_basis parameter.** Generally, DDTRN is not quite sensitive to num_basis variation and *num_basis* = 10 is adopted to show the prediction performance.

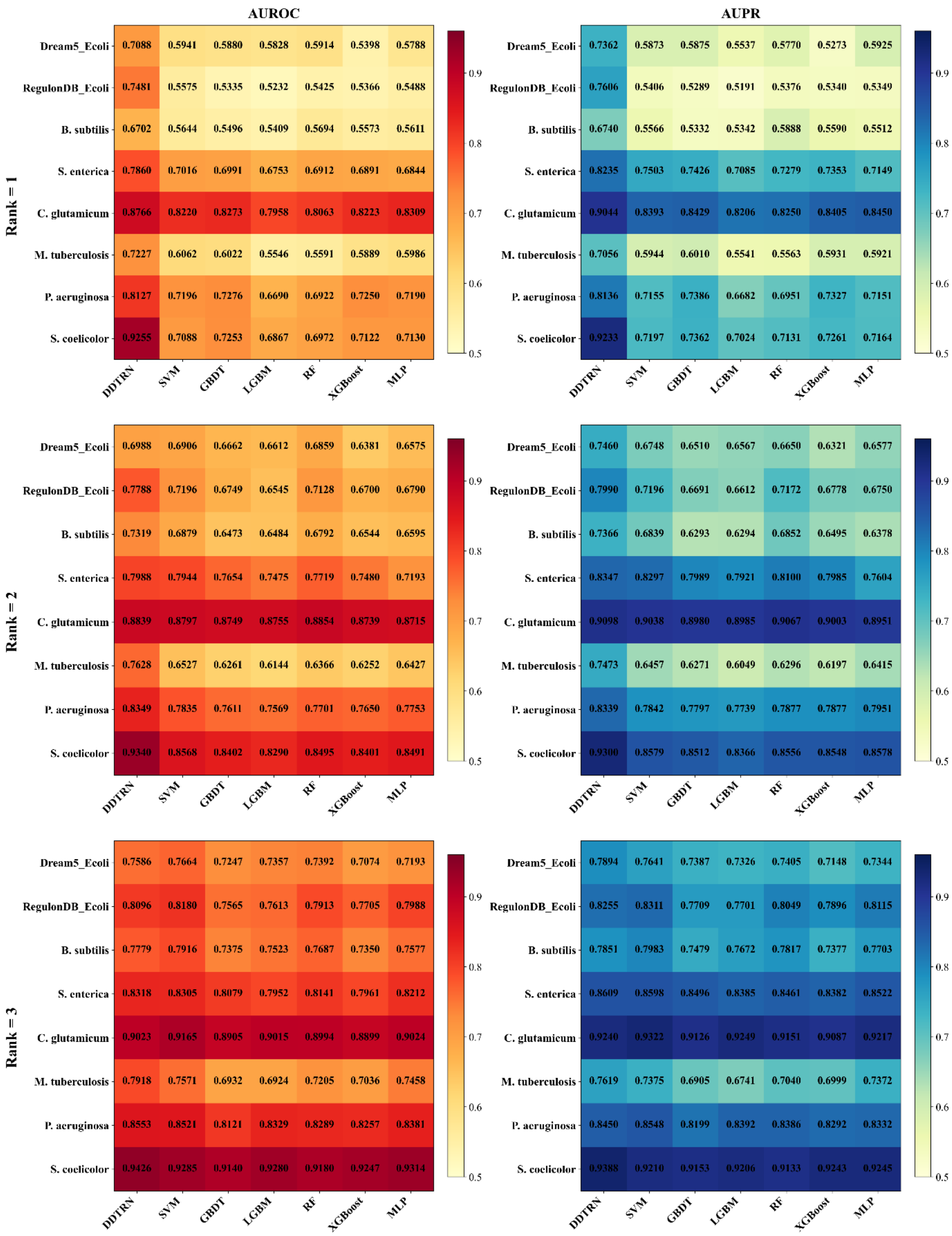
AUROC
AUPR
Rank = 1
Rank = 2
Rank = 3
DDTRN SVM GBDT LGBM RF XGBoost MLP
0.5 0.6 0.7 0.8 0.9
Rank = 1, AUROC:
Dream5_Ecoli 0.7088 0.5941 0.5880 0.5828 0.5914 0.5398 0.5788
RegulonDB_Ecoli 0.7481 0.5575 0.5335 0.5232 0.5425 0.5366 0.5488
B. subtilis 0.6702 0.5644 0.5496 0.5409 0.5694 0.5573 0.5611
S. enterica 0.7860 0.7016 0.6991 0.6753 0.6912 0.6891 0.6844
C. glutamicum 0.8766 0.8220 0.8273 0.7958 0.8063 0.8223 0.8309
M. tuberculosis 0.7227 0.6062 0.6022 0.5546 0.5591 0.5889 0.5986
P. aeruginosa 0.8127 0.7196 0.7276 0.6690 0.6922 0.7250 0.7190
S. coelicolor 0.9255 0.7088 0.7253 0.6867 0.6972 0.7122 0.7130
Rank = 1, AUPR:
Dream5_Ecoli 0.7362 0.5873 0.5875 0.5537 0.5770 0.5273 0.5925
RegulonDB_Ecoli 0.7606 0.5406 0.5289 0.5191 0.5376 0.5340 0.5349
B. subtilis 0.6740 0.5566 0.5332 0.5342 0.5888 0.5590 0.5512
S. enterica 0.8235 0.7503 0.7426 0.7085 0.7279 0.7353 0.7149
C. glutamicum 0.9044 0.8393 0.8429 0.8206 0.8250 0.8405 0.8450
M. tuberculosis 0.7056 0.5944 0.6010 0.5541 0.5563 0.5931 0.5921
P. aeruginosa 0.8136 0.7155 0.7386 0.6682 0.6951 0.7327 0.7151
S. coelicolor 0.9233 0.7197 0.7362 0.7024 0.7131 0.7261 0.7164
Rank = 2, AUROC:
Dream5_Ecoli 0.6988 0.6906 0.6662 0.6612 0.6859 0.6381 0.6575
RegulonDB_Ecoli 0.7788 0.7196 0.6749 0.6545 0.7128 0.6700 0.6790
B. subtilis 0.7319 0.6879 0.6473 0.6484 0.6792 0.6544 0.6595
S. enterica 0.7988 0.7944 0.7654 0.7475 0.7719 0.7480 0.7193
C. glutamicum 0.8839 0.8797 0.8749 0.8755 0.8854 0.8739 0.8715
M. tuberculosis 0.7628 0.6527 0.6261 0.6144 0.6366 0.6252 0.6427
P. aeruginosa 0.8349 0.7835 0.7611 0.7569 0.7701 0.7650 0.7753
S. coelicolor 0.9340 0.8568 0.8402 0.8290 0.8495 0.8401 0.8491
Rank = 2, AUPR:
Dream5_Ecoli 0.7460 0.6748 0.6510 0.6567 0.6650 0.6321 0.6577
RegulonDB_Ecoli 0.7990 0.7196 0.6691 0.6612 0.7172 0.6778 0.6750
B. subtilis 0.7366 0.6839 0.6293 0.6294 0.6852 0.6495 0.6378
S. enterica 0.8347 0.8297 0.7989 0.7921 0.8100 0.7985 0.7604
C. glutamicum 0.9098 0.9038 0.8980 0.8985 0.9067 0.9003 0.8951
M. tuberculosis 0.7473 0.6457 0.6271 0.6049 0.6296 0.6197 0.6415
P. aeruginosa 0.8339 0.7842 0.7797 0.7739 0.7877 0.7877 0.7951
S. coelicolor 0.9300 0.8579 0.8512 0.8366 0.8556 0.8548 0.8578
Rank = 3, AUROC:
Dream5_Ecoli 0.7586 0.7664 0.7247 0.7357 0.7392 0.7074 0.7193
RegulonDB_Ecoli 0.8096 0.8180 0.7565 0.7613 0.7913 0.7705 0.7988
B. subtilis 0.7779 0.7916 0.7375 0.7523 0.7687 0.7350 0.7577
S. enterica 0.8318 0.8305 0.8079 0.7952 0.8141 0.7961 0.8212
C. glutamicum 0.9023 0.9165 0.8905 0.9015 0.8994 0.8899 0.9024
M. tuberculosis 0.7918 0.7571 0.6932 0.6924 0.7205 0.7036 0.7458
P. aeruginosa 0.8553 0.8521 0.8121 0.8329 0.8289 0.8257 0.8381
S. coelicolor 0.9426 0.9285 0.9140 0.9280 0.9180 0.9247 0.9314
Rank = 3, AUPR:
Dream5_Ecoli 0.7894 0.7641 0.7387 0.7326 0.7405 0.7148 0.7344
RegulonDB_Ecoli 0.8255 0.8311 0.7709 0.7701 0.8049 0.7896 0.8115
B. subtilis 0.7851 0.7983 0.7479 0.7672 0.7817 0.7377 0.7703
S. enterica 0.8609 0.8598 0.8496 0.8385 0.8461 0.8382 0.8522
C. glutamicum 0.9240 0.9322 0.9126 0.9249 0.9151 0.9087 0.9217
M. tuberculosis 0.7619 0.7375 0.6905 0.6741 0.7040 0.6999 0.7372
P. aeruginosa 0.8450 0.8548 0.8199 0.8392 0.8386 0.8292 0.8332
S. coelicolor 0.9388 0.9210 0.9153 0.9206 0.9133 0.9243 0.9245

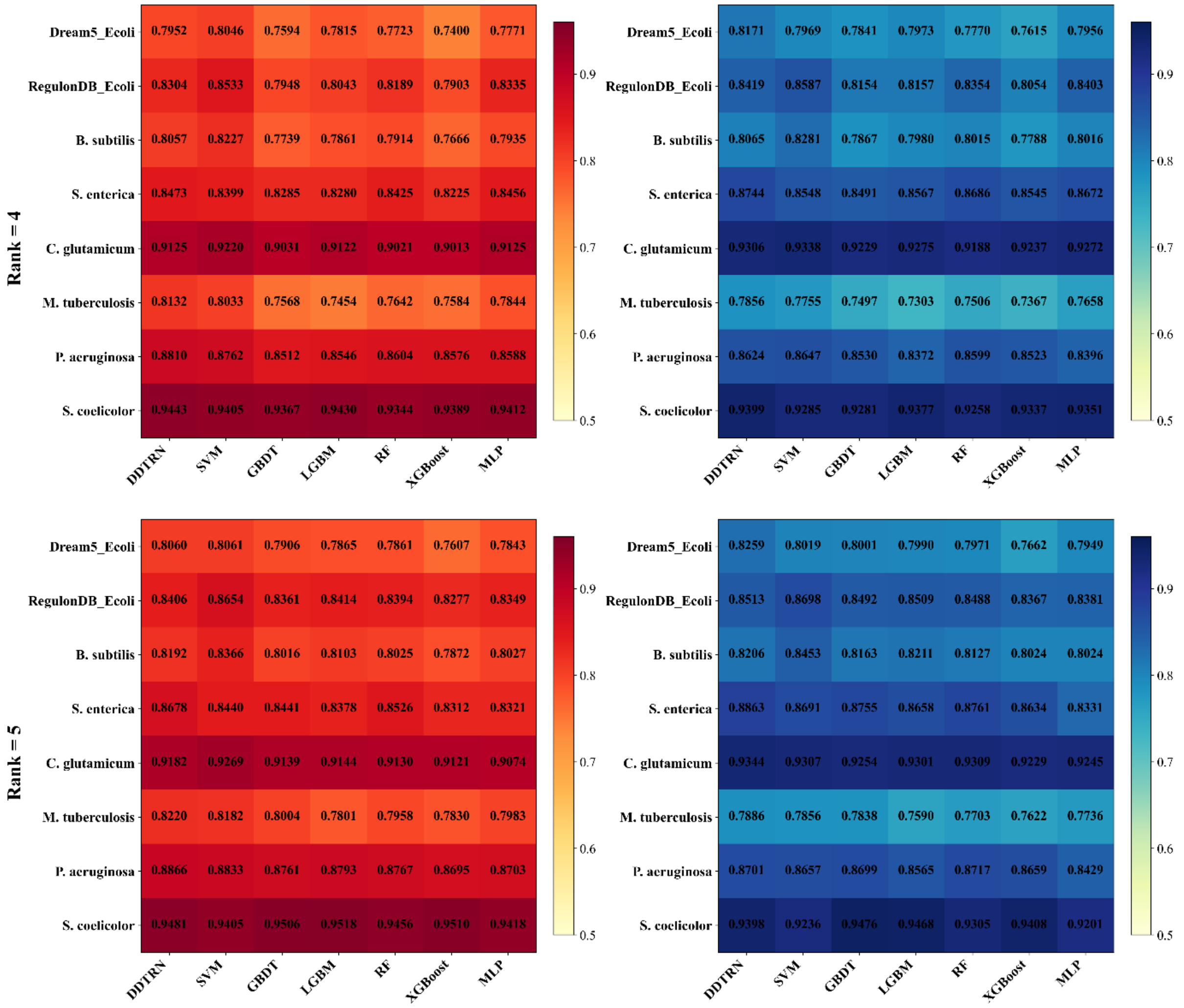


**Figure S3. Heatmaps showing the AUROC and AUPR scores of different methods on the benchmark datasets at ranks 1 to 5.** AUROC and AUPR are shown side by side for all methods across eight benchmark datasets. Color intensity increases with higher metric value.

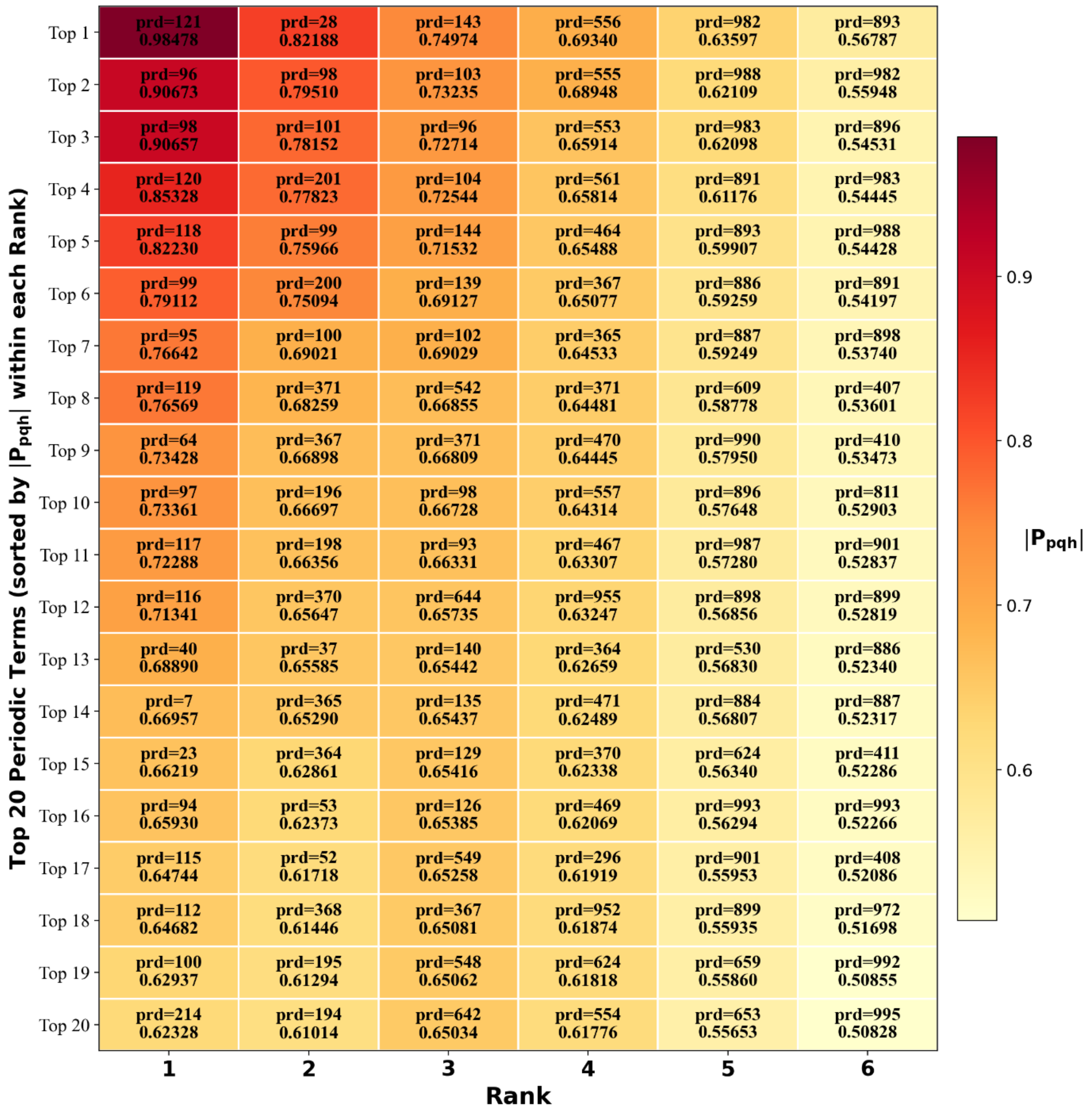


**Figure S4. Distribution of $|P_{pgh}|$ values for periodic terms in the PWF of DDTRN across different rank settings (Linear mode).** Heatmap shows the top 20 periodic terms sorted by the absolute value of the elements in tensor **P** (namely $|P_{pqh}|$) across ranks 1 to 6. “prd” indicates the corresponding periodic terms defined in Eq. (2). Larger $|P_{pqh}|$ value indicates greater contribution from the corresponding periodicity in the sequence pattern differentiating Regulatory and Non-regulatory interaction.
